# Large Quantities of Bacterial DNA and Protein in Common Dietary Protein Source Used in Microbiome Studies

**DOI:** 10.1101/2023.12.07.570621

**Authors:** Alexandria Bartlett, J. Alfredo Blakeley-Ruiz, Tanner Richie, Casey M. Theriot, Manuel Kleiner

**Author notes:** Correspondence: Manuel Kleiner –. Tanner Richie, Division of Biology, Kansas State University Manhattan, KS.

## Abstract

Diet has been shown to greatly impact the intestinal microbiota. To understand the role of individual dietary components, defined diets with purified components are frequently used in diet-microbiota studies. Many of the frequently used defined diets use purified casein as the protein source. Previous work indicated that this casein contains microbial DNA potentially impacting results of microbiome studies. Other diet-based microbially derived molecules that may impact microbiome measurements, such as proteins detected by metaproteomics, have not been determined for casein. Additionally, other protein sources used in microbiome studies have not been characterized for their microbial content. We used metagenomics and metaproteomics to identify and quantify microbial DNA and protein in a casein-based defined diet to better understand potential impacts on metagenomic and metaproteomic microbiome studies. We further tested six additional defined diets with purified protein sources with an integrated metagenomic-metaproteomic approach and show that contaminating microbial protein is unique to casein within the tested set as microbial protein was not identified in diets with other protein sources. We also illustrate the contribution of diet-derived microbial protein in diet-microbiota studies by metaproteomic analysis of stool samples from germ-free mice (GF) and mice with a conventional microbiota (CV) following consumption of diets with casein and non-casein protein. This study highlights a potentially confounding factor in diet-microbiota studies that must be considered through evaluation of the diet itself within a given study.

**Importance:** Many diets used in diet-microbiota studies use casein as the source of dietary protein. We found large quantities of microbial DNA and protein in casein-based diets. This microbial DNA and protein are resilient to digestion as it is present in fecal samples of mice consuming casein-based diets. This contribution of diet-derived microbial DNA and protein to microbiota measurements may influence results and conclusions and must therefore be considered in diet-microbiota studies. We tested additional dietary protein sources and did not detect microbial DNA or protein. Our findings highlight the necessity of evaluating diet samples in diet-microbiota studies to ensure that potential microbial content of the diet can be accounted for in microbiome measurements.

## OBSERVATION

The intestinal microbiota is highly influential to the health of the host (1, 2). Diet has been shown to shape the intestinal microbiota, yet we are still unraveling how individual dietary components impact the functioning of the microbiota (1, 3, 4). To understand the role of individual dietary components, many studies use purified dietary components, for example as a supplement in human feeding trials (5, 6) or as part of a defined diet in animal studies (1, 7). Diet-microbiota studies frequently use casein as the purified protein component of a defined diet, as casein is the protein source in the defined AIN-93 laboratory rodent diet (8).

Previous studies have shown that sterilized, purified casein protein used in defined diets contains significant amounts of microbial DNA. 16S rRNA gene sequencing identified the Gram-positive bacterium *Lactococcus lactis*, which is used in casein production, as the source of the microbial DNA (9-12). *Lactococcus*, and in some cases specifically *L. lactis*, have been identified by sequencing-based studies as a key microbiota member responding to a specific treatment, for example a high fat diet, in studies using casein (13, 14). While in some of these studies the dietary source of *L. lactis* was recognized (9-11), other studies do not show any indication of considering a potential dietary source of *Lactococcus*. In the studies that did recognize the potential issue with *L. lactis* contamination from the diet, two methods for addressing the issue have been used; bioinformatic removal of *L. lactis* reads during analysis (10, 12) or use of ethanol-washed casein in diets (12).

While the detection of *L. lactis* DNA in studies with casein-based diets has highlighted the importance of knowing the microbial content of dietary protein sources, our understanding of the breadth of the issue is limited. First, we currently only know about the presence of microbial DNA in casein, however, the presence of other microbially derived biomolecules such as protein could also critically impact microbiota measurements. Specifically, metaproteomics, which is used to study functional interactions in the microbiota by identifying and quantifying host and microbial proteins (15), would be impacted by microbial proteins introduced through the diet.

The contribution of diet-derived microbial protein to metaproteomic measurements of the microbiota has, however, not been previously investigated. Second, there is an increasing interest in studying the effect of different sources of dietary protein on the microbiota and resulting effects on host health (3). The microbial content of other dietary protein sources that are used in diet-microbiota studies has not yet been investigated.

### Massive quantities of microbial DNA and protein in purified casein diet

To assess the microbial protein content and its relevance in purified diets, we performed metagenomic sequencing and metaproteomics of 1) defined mouse diets with a single source of purified dietary protein (lactic casein or soy protein isolate), conventional (CV) and germ-free (GF) mouse stool samples following consumption of the two diets (Fig. 1A). The metagenomic sequencing allowed us to not only build a sample-matched protein sequence database for our metaproteomic analyses (16), but also to expand on previous reports which primarily relied on 16S rRNA gene sequencing and only investigated casein-based diets and casein-fed mice. We mapped the metagenomic reads to six reference genomes: *Mus musculus, Glycine max, Bos taurus, Thioflavicoccus mobilis* (control), *L. lactis* sequences retrieved from the assembled metagenome and the assembled metagenome with *L. lactis* sequences removed. While the reads from the soy diet mapped overwhelmingly to the soy genome, approximately 90% of reads from the casein diet mapped to the *L. lactis* reference (Fig. 1B).

**Figure 1.**
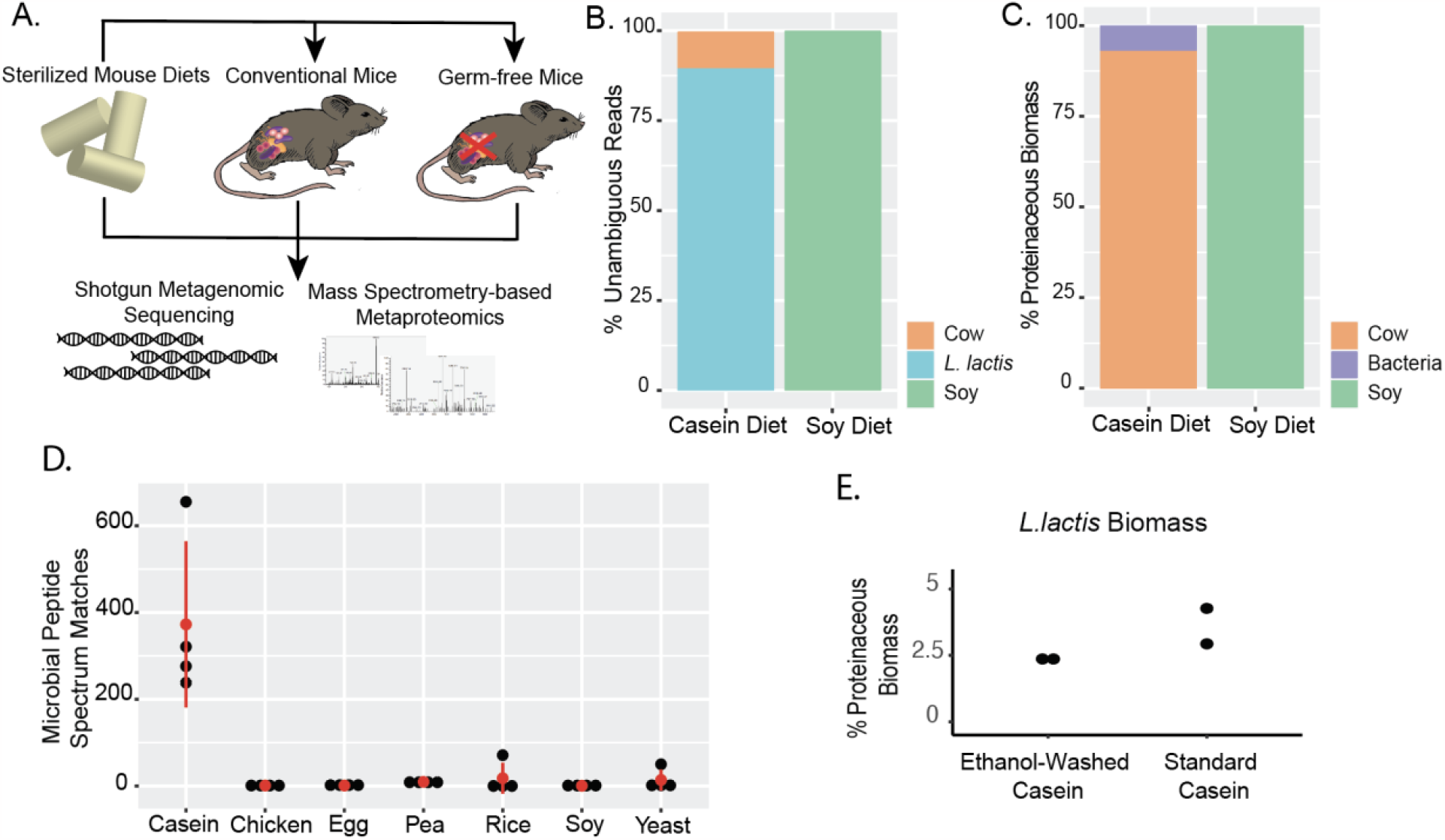
Experimental design and quantification of bacterial DNA and protein in diets. A) Metagenomic sequencing and mass spectrometry-based metaproteomics was performed on defined diets containing a single dietary protein source and on stool samples from germ-free mice and mice with a conventional microbiota. The stool samples were collected from the mice following consumption of diets containing purified soy protein or casein as the only protein source. The diets were sterilized prior to measurement and feeding using gamma irradiation. B) Average relative abundance of shotgun metagenomic reads from defined diet samples (n=3 per diet) that mapped unambiguously to diet reference genomes and assembled microbiota metagenomes. C) Proteinaceous biomass composition of sterilized diets (n=4 per diet) determined with metaproteomics according to the approach described by Kleiner et al (17). Bacterial proteins were identified using protein sequences predicted from the assembled shotgun metagenome to have a protein sequence database that matches the actual samples as described in Blakeley-Ruiz and Kleiner (16). Approximately 4.5% of the bacterial proteinaceous biomass in the casein diet samples was classified as *L. lactis* (Supplementary Table 3). The remaining 2.4% of the bacterial proteinaceous biomass in the casein diet samples were unbinned or unambiguous proteins in our microbiota database, which likely also represent *L*.*lactis* proteins. D) Number of bacterial peptide spectrum matches (PSMs) in various defined diets with 20% of a purified protein as the only protein source. The mean and standard deviation are indicated in red. E) Bacterial PSMs in ethanol-washed casein vs unwashed casein.

To quantify the microbial protein in the purified diets, we identified and quantified proteins in the soy (n=4) and casein (n=4) diets by LC-MS/MS and assessed the relative proteinaceous biomass (17). Soy protein represented the entirety of the protein in the soy diet but microbial proteins represented 4.7% of the proteinaceous biomass of the casein diet (Fig. 1C).

To investigate the microbial protein content of other dietary protein sources used in defined diets, we performed metaproteomic analysis on seven different defined (20% protein by weight) diets. In addition to the casein and soy diets described above, we analyzed diets formulated with alternative purified protein sources including Egg White Solids, Torula Yeast, Chicken Bone Broth, Yellow Pea, and Brown Rice (n=4 per diet). We observed more than 200 Peptide Spectrum Matches (PSMs) to bacterial proteins for all replicates for only the casein diet (Fig. 1D). We did observe 71 bacterial PSMs for one replicate of the rice diet and 50 bacterial PSMs for one replicate of the yeast diet, which we attributed to carryover from a sample with high bacterial content run on the LC-MS/MS system immediately prior to these two samples.

A previous study showed that ethanol-washed casein contains 1,000-fold less *L. lactis* DNA and suggested ethanol-washed casein as a potential alternative to the standard preparation of casein currently used in purified diets (12). To assess if ethanol-washed casein has a similar reduction in *L. lactis* protein, we assessed the proteinaceous biomass contribution of *L. lactis* in ethanol-washed casein and standard casein. Although the *L. lactis* protein content in ethanol-washed casein (2.4%) was on average lower than the standard casein (3.6%), the reduction was only minor and not comparable to the previously reported reduction in *L. lactis* DNA (Fig. 1E). This suggests that ethanol washing of casein is not a viable strategy for reducing *L. lactis* protein content of casein-based diets.

### Massive quantities of diet-derived microbial DNA and protein in stool of CV and GF mice fed casein diet

To examine the contribution of diet-derived microbial DNA and protein to metagenomic and metaproteomic measurements of the microbiota, we performed shotgun metagenomics and metaproteomics on stool collected from four cages of CV and GF mice after mice were fed the casein or soy diet for 7 days (Fig. 1A). We mapped the raw metagenomic reads to the 6 references described above. Over 24 million reads from the casein-fed GF mice mapped to the *L. lactis* reference (Fig. 2A). The majority of reads from the soy and casein CV mouse samples mapped to the Non-*L*.*lactis* microbiota reference, however, the casein CV mouse samples had an average of 1,291,589 reads that mapped to the *L. lactis* reference. In the metaproteomic analyses, we measured an average of 1,527 *L. lactis* PSMs in the casein-fed GF mice (Fig. 2B). As expected, the microbial PSMs we identified in the CV mice were primarily from Non-*L. Lactis* microbiota proteins (5,713 PSMs in casein-fed and 7,999 PSMs in soy-fed mice). However, in the casein-fed CV mice we measured on average, an additional 2,444 *L. Lactis* PSMs. Our data suggests that a large quantity of diet-derived *L. lactis* DNA and protein withstands the passage through the intestinal tract.

**Figure 2:**
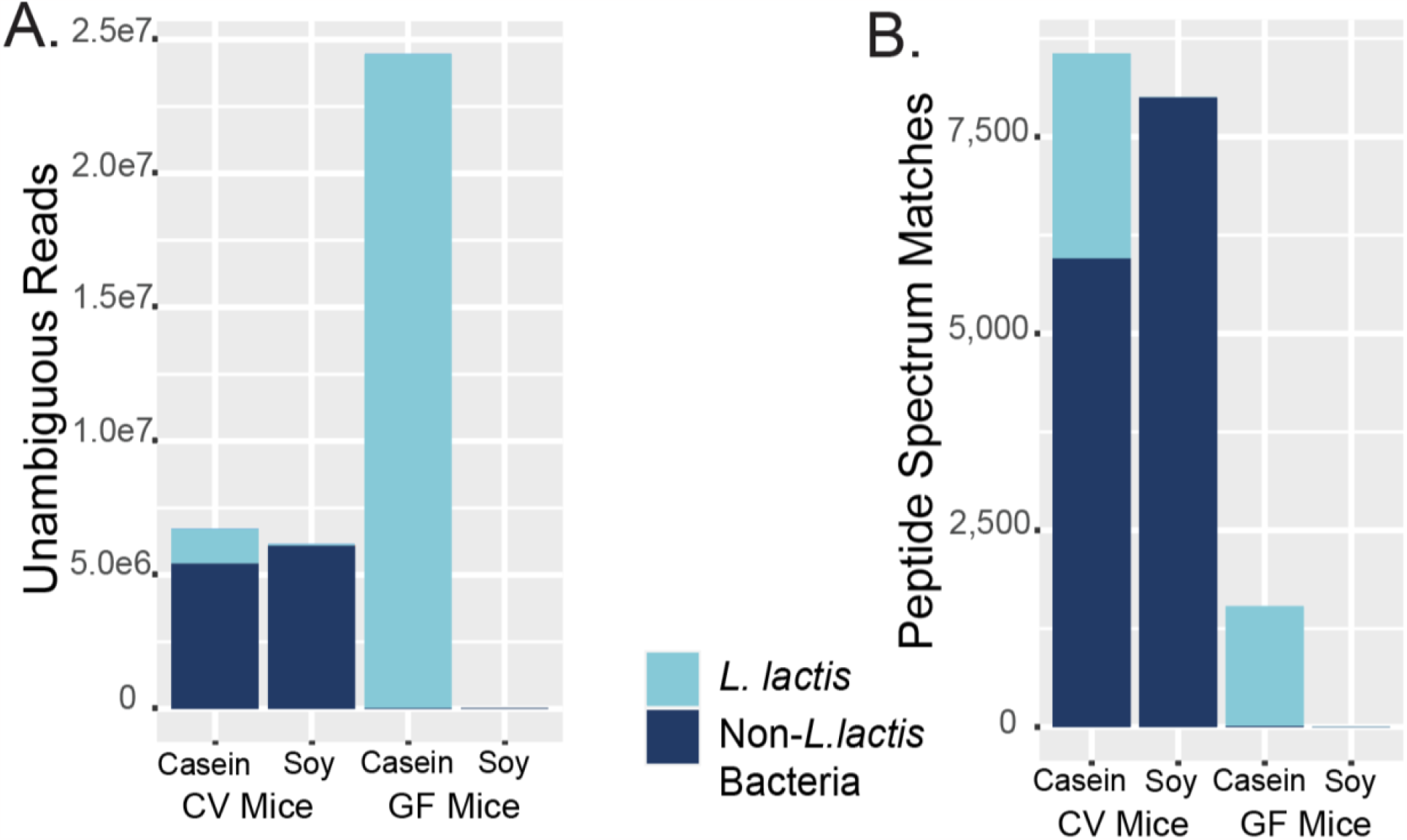
Quantification of bacterial proteins and DNA in fecal samples from germ-free (GF) mice and mice with a conventional microbiota (CV) following consumption of a defined diet with either purified soy protein or casein as the sole protein source. A) Average number of shotgun metagenomic reads from GF (n=3 per diet) and CV (n=3 per diet) mice fecal samples that mapped unambiguously to diet reference genomes and assembled microbiota metagenomes. B) Average microbial PSMs in GF (n=12) and CV mice (n=11-12).

## Conclusions

Here we show that one purified protein source (casein) used in diet-microbiota studies contains high amounts of microbial DNA and protein. This diet-derived microbial DNA and protein withstand passage through the intestinal tract and are thus present in metagenomic and metaproteomic measurements of the microbiota. Additionally, we show that a diversity of other purified dietary protein sources do not contain measurable amounts of microbial DNA and protein, based on the metagenomic and metaproteomic approaches used in this study. It is, however, to be expected that other purified dietary components (lipids, fiber, other protein sources etc.) and unpurified foods and drinks contain microbial DNA and protein to various extents, which may be present in microbiome measurements. In particular, foods and drinks that involve microbial fermentation in their preparation such as, bread, cheese, yogurt, kimchi and beer may contain significant amounts of microbial DNA (e.g. Fig. 4 in (4)) and protein, even if the microbes from which the DNA and protein originate are non-viable in the intestinal tract.

Therefore, in any study that seeks to evaluate the impact of diet on the microbiota, the content of microbial compounds in the diet that might confound the measurements should be evaluated using the same measurement approach as the one applied to the intestinal or fecal samples. The obtained information on the microbial content of dietary components can then be used to either choose diets with a lower amount of microbial compounds or alternatively to bioinformatically remove/account for the known microbial compounds from the diet in the data obtained from the intestinal/fecal sample measurements.

### NCBI bioproject, ProteomeXchange Consortium and Dryad Data Repository

Sequencing data available under bioprojects PRJNA1026909 (microbial database) and PRJNA1026974 (metagenomic dataset). Mass spectrometry data and protein sequence databases available via PRIDE repository under PXD041586 and PXD040649. Proteins identified at 5% FDR in diet and mouse samples available via Dryad data repository with DOI: 10.5061/dryad.nvx0k6dzq. See data accessibility in materials and methods for details.

## Materials and Methods

### Animals and housing

12 conventional C57BL/J6 mice (6 males, 6 females, Jackson Labs Bar Harbor) and 12 germ-free C57BL/J6 mice (6 males, 6 females, NCSU gnotobiotic core) were used in this study. All mice were 3-6 months in age and housed in groups of two or three by sex. The food, bedding and water were autoclaved. All conventional mouse cage changes were performed in a laminar flow hood. The mice were subjected to a 12 h light and 12 h dark cycle and were housed at an average temperature of 70F and 35% humidity. Animal experiments were conducted in the Laboratory Animal Facilities located on the NCSU CVM campus. The animal facilities are managed by full-time animal care staff coordinated by the Laboratory Animal Resources (LAR) division at NCSU. The NCSU CVM is accredited by the Association for the Assessment and Accreditation of Laboratory Animal Care International (AAALAC). Trained animal handlers in the facility fed and assessed the status of animals several times per day. Those assessed as moribund were humanely euthanized by CO2 asphyxiation. This protocol is approved by NC State’s Institutional Animal Care and Use Committee (IACUC).

### Animal diets and sample collection

All diets used in this study were irradiated, not supplemented with amino acids and contained a single source of dietary protein (Supplementary Table 1). The 20% soy diet (Envigo Teklad Diets) was fed *ad libitum* to both conventional and germ-free mice for 7 days. After 7 days of the soy diet, fecal samples were collected and the soy diet was replaced with a 20% casein diet.

After 7 days of the casein diet, fecal samples were collected again. Fecal samples were collected into NAP buffer preservation solution at a ratio of approximately 1:10 Sample Weight: Preservation Solution Volume, and roughly homogenized with a sterilized disposable pestle (18-20). All animal protocols were approved by the Institutional Animal Care and Use Committee of North Carolina State University.

### Metagenomic sequencing

Two different shotgun metagenomic datasets were generated in this study. The first round of sequencing was done to prepare a matched metagenomic-based metaproteomic database for the identification of microbiota proteins in the CV mice (PRJNA1026909). In the methods described below, this dataset is referred to as the microbial database. The microbial database was used 1) to identify the microbial proteins via metaproteomic analysis and 2) as a reference for the read mapping of the second metagenomic dataset. The second round of metagenomic sequencing was conducted to identify the DNA present in the diet samples, GF mice and CV mice (PRJNA1026974). This dataset is referred to as the metagenomic dataset.

### DNA extraction for the Microbial Database

To generate a matched metagenomic-based proteomic database of the conventional mice microbiota, DNA was extracted from 16 different CV mouse samples. These 16 samples represented multiple sampling points from four different cages of conventional mice, collected 7 days after mice consumed different purified protein diets. The protocol described by Knudsen *et al*., which is based on the QIAamp DNA stool mini kit (Qiagen), was used to extract DNA, with minor modifications (21). To remove the preservation solution from the fecal samples, 5 mL of 1X Phosphate Buffered Saline solution (VWR) was added to samples to dilute the preservation solution, followed by centrifugation (17,000 x g, 5 min) to pellet solids and bacterial cells.

Pellets were lysed by beadbeating (3.1 m/s for 3 cycles of 30 sec. with 1 min of cooling on ice in between each cycle) in 2 ml bead beating tubes (Lysing Matrix E, MP Biomedicals) using a Bead Ruptor Elite 24 (Omni International). DNA concentration of eluates was assessed using a DS-11 FX+ Spectrophotometer (Denovix) using a Qubit™ dsDNA High Sensitivity Assay Kit (Invitrogen). 200 ng of each individually extracted sample was used to pool by cage (4 samples sent for sequencing).

### DNA extraction for the Metagenomic Dataset

DNA was extracted from fecal samples of 3 GF mice and 3 CV mice (2 male, 1 female) both after feeding on the soy diet and the casein diet (12 samples in total), 3 replicates of the irradiated soy diet, 3 replicates of the casein diet and a *Thioflavicoccus mobilis* (*T. mobilis*) culture as a control. We used the same protocol for DNA extraction as above.

### DNA sequencing for the Microbial Database

Metagenomic DNA was submitted to the North Carolina State Genomic Sciences Laboratory (Raleigh, NC, USA) for Illumina library construction and sequencing to produce between 51,152,549 and 74,618,259 150 bp paired-end reads for each of the 4 samples. Library construction was performed using an Illumina TruSeq Nano Library kit according to manufacturer’s instructions. Libraries were sequenced on an Illumina NovaSeq 6000 sequencer.

### DNA sequencing for the Metagenomic Dataset

Metagenomic DNA was submitted to Diversigen for Illumina library preparation and sequencing using a single lane of a NovaSeq to generate 100 bp single-end reads. To mitigate index hopping, dual indexing was performed and a control sample was included (DNA extracted from *T. mobilis)* for assessment between samples in a single lane. The number of reads that unambiguously mapped to the *T. mobilis* reference was less than 0.25% of all unambiguously mapped reads for diet and mouse samples (Supplementary Table 2).

### Read processing, assembly, binning and annotation (Microbial Database)

The BBSplit algorithm was used to remove phix174 (NCBI GenBank accession CP004084.1) and mouse genome (mm10) reads, followed by quality trimming with BBDuk (BBMap, Version 38.06) using the following settings: mink = 6, minlength =20. MetaSPAdes (version 3.12.0) with error correction and k-mer lengths 33, 55, 99 was used to assemble the 4 sequenced samples individually (22). In addition to assembling the samples individually, a co-assembly of the four samples was performed using MEGAHIT (Version 1.2.4) to increase the number of high-quality Metagenome Assembled Genomes (MAGs). MetaBAT (version 2.12.1) was used to bin the assembled contigs and the resulting MAGs were evaluated using CheckM (version 1.1.2) (23, 24). MAGs with a CheckM quality score of >50 completeness <10 were considered medium quality and accepted for further consideration. MAGs with >30 completeness and <5 contamination were also included to avoid missing small genomes or taxonomic groups that did not assemble well. dRep (Version 2.6.2) was used to cluster the MAGs into species groups at 95% average nucleotide identity (25). PROKKA (Version 1.14.6) was used for gene prediction of the MAGS and unbinned contigs. Taxonomy of the MAGs was predicted using GTDB-Tk (Version 1.3.0) using reference database r95 and BAT(Version 5.0.3) (26, 27).

### Protein sequence database construction for metaproteomics

A non-redundant microbiota protein sequence database was constructed with the annotated protein sequences from the microbial database to identify microbial proteins via metaproteomic analysis. To remove redundant protein sequences, protein sequences from MAGs were clustered with an identity threshold of 95% using cd-hit (Version 4.7) (28). Protein sequences from unbinned contigs were separately clustered at 95% similarity. Cd-hit-2d was used to identify sequences from unbinned contigs and low-quality MAGs that were not represented in the set of binned sequences with at least 90% similarity. Sequences with less than 90% similarity to binned sequences were added to the microbiota protein sequence database.

The microbiota protein sequence database was combined with the *Mus musculus* reference proteome (UP000000589, Downloaded 19Feb20) and one of the respective dietary proteomes to generate six different databases. The dietary reference proteomes included *Glycine max* (UP000008827, Downloaded 19Feb20), *Bos taurus* (UP000009136, Downloaded 19Feb20), *Cyberlindnera jadinii* (UP000094389, Downloaded 25May20), *Oryza sativa* (UP000059680, Downloaded 25May20) and *Gallus gallus* (UP000000539, Downloaded 25May20). Due to the lack of a reference proteome for the yellow pea diet, we created a custom pea reference with all available UniProtKB protein sequences for *Pisum sativum* (Taxon ID: 388 Downloaded 25Apr20) and the reference proteome of *Cajanus cajan* (UP000075243, Downloaded 25May20). Each reference proteome and the *Pisum sativum* protein sequences were clustered individually with an identity threshold of 95% using cd-hit (28).

### Read mapping of metagenomic dataset

The BBsplit algorithm was used to map raw reads from the metagenomic dataset to six references. The references consisted of the *Mus musculus* genome (GCA_000001635.9), *Glycine max* genome (GCA_000004515.4), *Bos taurus* genome (GCF_002263795.1), *T. mobilis* (GCF_000327045.1), *L. lactis* and Non-*L. lactis* microbiota. The *L. lactis* reference comprised the seven MAGs from our microbial dataset that were taxonomically classified as *L. lacti*s. The Non-*L. lactis* microbiota reference consisted of all other high-quality MAGs not classified as *L. lactis* from our metagenomic database. The following BBsplit parameters were used: ambiguous2=toss, qtrim=lr, minid=0.97. Plots were made using ggplot2 (Version 3.4.0) in Rstudio (Version 4.1.1) (29, 30).

### Protein extraction, peptide preparation and determination of diet and fecal samples

We removed the NAP buffer preservation solution from fecal samples by centrifugation (21,000 x g, 5 min). As diet samples were not collected in preservation solution, 100 mg was placed directly into Lysing Matrix E tubes (MP Biomedicals) for each replicate. Cells were lysed and proteins solubilized with SDT lysis buffer [4% (w/v) SDS, 100 mM Tris-HCl pH 7.6, 0.1 M DTT] and bead beating in Lysing Matrix E tubes (MP Biomedicals) (5 cycles of 45s at 6.45 m/s, 1 min between cycles). Following bead beating, samples were heated to 95°C for 10 min., then centrifuged (21,000 x g, 5 min). Tryptic digests were prepared (16 hour digestion) using the filter-aided sample preparation protocol (31). In brief, 60 µl of lysate was combined with 400 µl of UA solution (8 M urea in 0.1 M Tris/HCl pH 8.5) in 10 kDa MWCO 500 µl centrifugal filters (VWR International). Samples were centrifuged at 14,000 *g* for 30 min. Depending on sample concentration, this step was repeated up to three times to load the filter to capacity. Filters were washed with 200 µl of UA solution at 14,000 *g* for 40 min. 100 µl IAA (0.05 M iodoacetamide in UA solution) was added to filters, incubated for 20 min, and centrifuged at 14,000 *g* for 20 min.

Filters were washed with 100 µl of UA three times, followed by three washes of 100 µl ABC (50 mM Ammonium Bicarbonate). For digestion, 0.95 µg of MS grade trypsin (Thermo Scientific Pierce, Rockford, IL, USA) in 40 µl of ABC was added to filters and incubated for 16 hours in a wet chamber at 37 °C. Following digestion, samples were centrifuged at 14,000 *g* for 20 min. To elute peptides, 50 µl of 0.5 M NaCl was added and samples were centrifuged for 20 min. Peptide concentrations were determined with the Pierce Micro BCA assay (Thermo Scientific Pierce) according to the manufacturer’s instructions.

### LC-MS/MS of diet and fecal samples

We analyzed the peptides from fecal and diet samples by LC-MS/MS as previously described with small modifications (18). The samples were blocked and randomized to control for batch effects as previously described (32). An UltiMate™ 3000 RSLCnano Liquid Chromatograph (Thermo Fisher Scientific) was used to load peptides (600 ng for mouse fecal samples, 300 ng for diet samples) onto a 5 mm, 300 μm ID C18 Acclaim® PepMap100 pre-column (Thermo Fisher Scientific) with loading solvent A (2% acetonitrile, 0.05% TFA).

Peptides were then separated on an EASY-Spray analytical column heated to 60°C (PepMap RSLC C18, 2 µm material, 75 cm× 75 µm, Thermo Fisher Scientific) using a 140 min gradient at a flow rate of 300 nl/min. The first 102 minutes of the gradient went from 95% eluent A (0.1% formic acid) to 31% eluent B (0.1% formic acid, 80% acetonitrile), followed by 18 min from 31 to 50% B, and 20 min at 99% B. To reduce carryover, a wash run with 100% acetonitrile was inserted between samples. Eluting peptides were ionized by electrospray ionization and analyzed in a Q Exactive HF hybrid quadrupole-Orbitrap mass spectrometer (Thermo Fisher Scientific) with the following parameters: m/z 445.12003 lock mass, normalized collision energy equal to 24, 25 s dynamic exclusion, and exclusion of ions of +1 charge state. A full MS scan from 380 to 1600 *m*/*z* was performed at a resolution of 60,000 and a max IT of 200 ms. Data-dependent MS^2^ was performed for the 15 most abundant ions at a resolution of 15,000 and max IT of 100 ms.

### Protein identification and analysis of diet and fecal samples

A protein sequence database containing the matched metagenomic database and multiple reference proteomes was used to search the MS^2^ spectra. The Proteome Discoverer software version 2.3 (Thermo Fisher Scientific) was used for protein identification using run calibration and the Sequest HT node, with the following settings: trypsin (Full), maximum 2 missed cleavages, 10 ppm precursor mass tolerance, 0.1 Da fragment mass tolerance and maximum 3 equal dynamic modifications per peptide. The following dynamic modifications were considered: oxidation on M (+15.995 Da), deamidation on N,Q,R (0.984 Da) and acetyl on the protein N terminus (+42.011 Da). The static modification carbamidomethyl on C (+57.021 Da) was also included. The percolator node in Proteome Discoverer was used to calculate peptide false discovery rate (FDR) with the following parameters: maximum Delta Cn 0.05, a strict target FDR of 0.01, a relaxed target FDR of 0.05 and validation based on q-value. The Protein-FDR Validator node in Proteome Discoverer was used for protein inference with a strict target FDR of 0.01 and a relaxed target FDR of 0.05 to restrict protein FDR to below 5%.

To assess the contribution of microbial proteins in the diet and mouse samples, identified proteins were filtered for 5% FDR and 2 protein unique peptides and summed by organism, as described in Kleiner et al (17). Only organisms with at least 10 PSMs for all 4 replicates of each diet sample were included in the biomass assessment. Plots were made using ggplot2 (Version 3.4.0) in Rstudio (Version 4.1.1) (29, 30).

### Comparison of ethanol washed casein with standard lactic casein

Envigo gifted us two lots of ethanol washed vitamin-free casein and standard casein used in their purified protein diets. Proteins were extracted from 250 µg of sample and tryptic digests were prepared using the filter-aided sample preparation protocol described above (31). LC-MS/MS analysis was similar to the method described above for diet and fecal samples the only modification being that 400 ng of peptides were loaded onto the analytical column. To identify peptides and proteins, MS/MS spectra were searched against the same protein sequence database as described above. Proteins were considered *Lactococcus lactis* if they 1) were present in the *Lactococcus lactis* annotated MAGs from the matched metagenomic database or 2) matched to the Uniprot *Lactococcus lactis* reference proteomes UP000002196 and UP000015854 at a 95% identity threshold using diamond blastp (33). We then calculated percent proteinaceous biomass as described in Kleiner et al (17).

## Data accessibility

All sequencing data has been submitted to NCBI as part of bioprojects PRJNA1026909 (reviewer link: https://dataview.ncbi.nlm.nih.gov/object/PRJNA1026909?reviewer=a6ipv6iees1fq136cqt5rmq2q 4) and PRJNA1026974 (reviewer link: https://dataview.ncbi.nlm.nih.gov/object/PRJNA1026974?reviewer=t94m1sut42r15j9himabksubhp). The mass spectrometry data and protein sequence databases were deposited to the ProteomeXchange Consortium via the PRIDE (34) partner repository with the data set identifier PXD041586 [reviewer access at this link https://www.ebi.ac.uk/training/user/login, with credentials user, password:V9Jz2n4h] and PXD040649 [reviewer access at this link https://www.ebi.ac.uk/training/user/login, with credentials user, password:ucnMkbYg]. All proteins identified at 5% FDR in diet and mouse samples have been submitted to the Dryad data repository with DOI: 10.5061/dryad.nvx0k6dzq.

## Supplemental Material

Supplementary Table 1: Composition of diets used in this study.

Supplementary Table 2: Raw read numbers and unambiguously mapped read numbers from metagenomic dataset samples, including 3 replicates of soy diet, casein diet, GF and CV mice.

## 3.7 Acknowledgements

We are grateful to Fernanda Salvato for providing extensive training for the metaproteomic analysis, Alissa Rivera for assistance with conventional mouse sample collection, Karen Flores for technical assistance with manipulations of germ-free mice, Angie Mordant for assistance with mouse experiments and Susan Tonkonogy for guidance on germ-free mouse experiments. We thank Envigo for the kind gift of ethanol-washed casein samples. We thank the Genomic Sciences Laboratory at North Carolina State University for performing Metagenomic sequencing and the Bioinformatic and Statistical Consulting Cores at North Carolina State University for performing statistical analysis and consultation. All LC-MS/MS measurements were made in the Molecular Education, Technology, and Research Innovation Center (METRIC) at North Carolina State University. The Gnotobiotic Core at the College of Veterinary Medicine, North Carolina State University is supported by the National Institutes of Health funded Center for Gastrointestinal Biology and Disease, NIH-NIDDK P30 DK034987. This work was supported by the Foundation for Food and Agriculture Research Grant ID: 593607 and by the National Institute Of General Medical Sciences of the National Institutes of Health under Award Number R35GM138362.

